# Unravelling the mechanisms of adaptation to high pressure in proteins

**DOI:** 10.1101/2022.04.25.489375

**Authors:** Antonino Caliò, Michael Marek Koza, Stephane Fontanay, Philippe Oger, Judith Peters

## Abstract

Life is thought to have appeared in the depth of the sea, under high hydrostatic pressure. Nowadays, it is known that the deep biosphere hosts a myriad of life forms thriving under high pressure conditions. However, the evolutionary mechanisms leading to their adaptation are still not known. Here we show the molecular bases of these mechanisms through a neutron scattering study of two orthologous proteins. We observed that pressure adaptation involves the decoupling of protein-water dynamics and the elimination of cavities in the protein core. This is achieved by an enrichment of acidic residues on the protein surface and by the use of bulkier hydrophobic residues in the core. These findings will be the starting point in the search of a complete genomic model explaining high pressure adaptation.

## Introduction

According to one of the most credited hypotheses on the origins of life, it appeared in the deep sea^1^, protected from the deleterious radiation from the young sun, but close to energy sources (e.g. hydrothermal vents) that could sustain relevant chemical reactions^2^. Therefore, it would have appeared under high hydrostatic pressure (HHP) conditions. Modern HHP-adapted organisms (piezophiles) display a pressure-dependent physiology, however, deciphering their adaptation to HHP is a very challenging task, because it is often concomitant with other environmental adaptations^3^. Indeed they usually thrive in very cold^4^, i.e. the deep ocean, or very hot environments^5^, i.e. hydrothermal vents. The first demonstration of proteome structural adaptation in piezophiles originated from comparative whole cell studies between two nearly isogenic piezophilic and piezosensitive microorganisms, namely *T. barophilus* and *T. kodakarensis*, which share identical growth characteristics, except for HHP-adaptation. Their proteomes exhibited different dynamical properties, together with a remarkable difference in the response to HHP of the hydration water^6^. In contrast to accepted models, proteins of the piezosensitive microorganism appear less sensitive to increasing HHP, while those of the piezophile are more flexible, and undergo pressure-dependent rearrangements at a pressure value close to the optimum of the organism^6,7^. Above this threshold, the piezophile proteome becomes pressure-insensitive^6–9^. Thus, unexpectedly, the adaptation to HHP in piezophiles seems to imply that the cell’s proteome is both more sensitive and more resistant to HHP^6^. These dynamical characteristics can also be preserved by piezophile cells under low pressure stress, through the accumulation of organic osmolytes^10^. Interestingly, a similar insensitivity to HHP has been observed for concentrated protein solutions in presence of organic osmolytes^11^. Another peculiarity of piezophiles is the response of their proteome’s hydration shell to HHP: its size is reduced and water appears less mobile^6^. It is thus probable that structural adaptation in proteins of piezophiles affect amino-acids at the water-protein interface. Therefore, two different processes seem to be responsible for HHP adaptation: i) a structural (i.e. genomic) adaptation, modifying protein sequences to alter their dynamics, and/or ii) the modulation of protein-water interaction. To date, all attempts to identify the structural signature of HHP adaptation at the genome level has failed, likely because it only involves amino-acids that interact with the hydration water or take part in the formation of internal cavities, which can be greatly destabilized by HHP^12^. If the macroscopic thermodynamics of proteins under pressure is quite well established^12–14^, its influence on their microscopical properties and their dynamics is still a debated subject^15–21^. Many different, and often contrasting, contributions govern the structural and dynamical stability of proteins with respect to HHP^14^, such as the presence of solvent inaccessible cavities^12^, electrostriction^22^ and the pressure dependence of the hydrophobic effect^23^. Concerning the fast dynamics, there is evidence that pressure tends to slow it down and inhibit conformational changes that require large amplitude motions^15,21^. To investigate protein HHP adaptation, Elastic Incoherent Neutron Scattering (EINS) and Quasi-Elastic Neutron Scattering (QENS) have been employed to study the dynamics of two orthologous proteins from *T. barophilus* (Tba) and *T. kodakarensis* (Tko), which only differ by their optimal growth pressure. Our data show profound differences both in their dynamics and in their interaction with the surrounding water layer, giving the first hints about the molecular mechanisms involved in HHP adaptation.

## Results

### Protein characterization

Genes TERMP_00744 and TK_0503, coding for a *Phosphomannose isomerase* (PMI), were cloned into the pET-16b over-expression vector^24^. The two PMIs have been over-expressed in *E. coli* (BL21 (DE3) pLysS) and purified by heat-treating the cell lysate and by size-exclusion chromatography (fig. S3, see Methods for further details). The PMIs are dimers (fig. S3). Homology modelling was performed to obtain a model structure for each protein (fig. S5) using i-Tasser^25^ or Swiss-Model^26^. Both models present very similar features: the protein structure is dominated by *β*-sheet contributions with turns and disordered regions connecting them, which forms a jelly-roll barrel structure, and is correctly predicted to be dimeric (fig. S6). The putative active site is located inside the barrel (fig. S7), and is conserved between the two proteins (His44, His46, Glu51 and His85), as are the residues involved in the monomer-monomer contact, except two conservative substitutions (A17L and I100V). The two orthologs differ at 16 positions (fig. S1) and most of the substitutions are located at the protein water interface and involve mainly polar and charged residues (fig. S8), with the notable exceptions of F8L and F105H which are located at the ligand pocket entrance (fig. S9), and I35V, located in the hydrophobic core of the protein.

### Elastic incoherent neutron scattering (EINS)

EINS was used to access the motions of the two proteins^27,28^, and to probe their response to HHP, while considering their structural differences, in order to shed light on their adaptation strategies. Incoherent neutron scattering probes the single-particle self correlation function^29^ and in protein samples the signal is dominated by hydrogen atoms, thanks to their very large incoherent cross-section^30^. In the case of elastic scattering, there is no energy exchange between the incident neutrons and the sample meaning that, in the time domain, the correlation function is probed in the long-time limit. Given that the instrument has a finite energy resolution (8 *μeV* for IN13, see Methods), this time limit is not at infinity, but it defines the *time window* of the instrument (∼ 100 *ps* for IN13). Figure 1 shows the scattering curves for Tba PMI at three representative temperatures. As expected, the scattering intensity shows a general decrease with temperature, consistent with the activation of anharmonic motions^31^.

**Figure 1:**
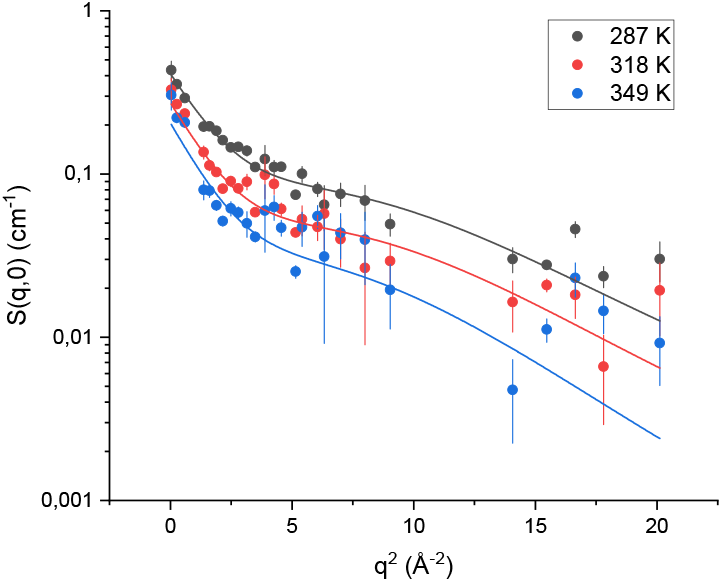
Scattering curves for Tba PMI at 150 bar and at some representative temperatures, with the corresponding two-state model fits.

Some contributions from global diffusion of the protein are likely present given the low concentration of the samples. Nonetheless, due to the very close similarity in the primary sequence and the almost identical molecular weight of the two proteins, it is reasonable to assume that these contributions would affect the measured signal in a similar fashion for both samples. Hence the observed differences can be ascribed to the distinct internal dynamics of the two proteins. Data have been interpreted in the framework of a two-state model^32^, which assumes two harmonic potential wells with an associated Mean Square Displacement (MSD) 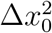, separated by a distance *d* and a free energy Δ*G* = Δ*H* − *T*Δ*S* (see Methods). The temperature-independence of *d* (fig. S10), Δ*H* and Δ*S* (consistently with the assumed Arrhenius behaviour of the two wells’ populations), allowed the employment a global fitting procedure at each pressure point, in which the only temperature-dependent parameter was 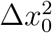. This granted the minimization of the number of free fitting parameters and greatly improved the quality and stability of the fittings. Figure 2 shows, for both samples, the single-well MSD, 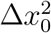, and the total MSD, 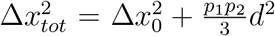, which takes into account the jump distance between the two wells and their populations. The absolute values of 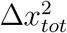, which report on the amplitude of the internal motions, appear very similar for both samples around 350*K*, indicating that the proteins display a similar degree of flexibility. However, the two samples show a clearly different temperature dependence of the MSD, i.e. the slope of the curve, inversely proportional to the protein’s resilience^33,34^: while Tko PMI displays a smooth increase, a change of slope in Tba PMI at around 320 *K* and 1 bar evidences the lower resilience, i.e. higher *softness*, of the protein.This transition is not present at higher pressures, and a linear temperature dependence of the MSD is observed. These results are in line with those from whole-cell studies on the same two organisms, which demonstrated the higher flexibility of the proteome of *T. barophilus*^6,7,35^ and the existence of pressure-induced structural rearrangements^7,35^ in this strain. For *T. barophilus*, the transition occurs smoothly between 1 and 300 bars, close to the optimal growth pressure for the organism, i.e. 400 bars^36^. In *T. kodakarensis*, the transition takes place at a much lower pressure range as shown by the sharp decrease of the MSD from 1 to 150 bars. Further striking differences between the two proteins can be found looking at the pressure dependence of the other parameters extracted from the two-state model fitting (fig. 3). In particular, the distance between the wells, *d* (panel a), is essentially pressure-independent for Tba PMI, while a sizeable decrease is detected for Tko PMI. This behaviour has already been observed in the model protein myoglobin^21^, and has been explained in terms of an increased roughness of the protein energy landscape, arising from the difficulty of the protein to explore the conformational substates characterized by bigger volume differences, in agreement with Le Châtelier’s principle^37^. In contrast, the energy landscape of Tba PMI appears to be extremely stable with respect to pressure application, resembling what Shrestha et al^20^ found for the *T. thioreducens* IPPase. However, in this case the comparison with the piezosensitive counterpart was considerably less significative, as hen egg white lysozyme is not related to the IPPase. The stability of Tba PMI is confirmed by the behaviour of the thermodynamic parameters Δ*H* (fig. 3b) and Δ*S* (fig. 3c), as they both decrease sharply from 1 to 150 bar and then become pressure-independent, while they both increase for Tko PMI. A decrease in the Δ*S* has been connected with decreased hydration^32^ and, indeed, such a decrease of the hydration shell size has been detected in the piezophilic proteome^6^. This is consistent with the two proteins having a different interaction with water. The exclusion of water from the hydration shell of Tba PMI appears to be the key of the dynamical stability of the protein under high pressure.

**Figure 2:**
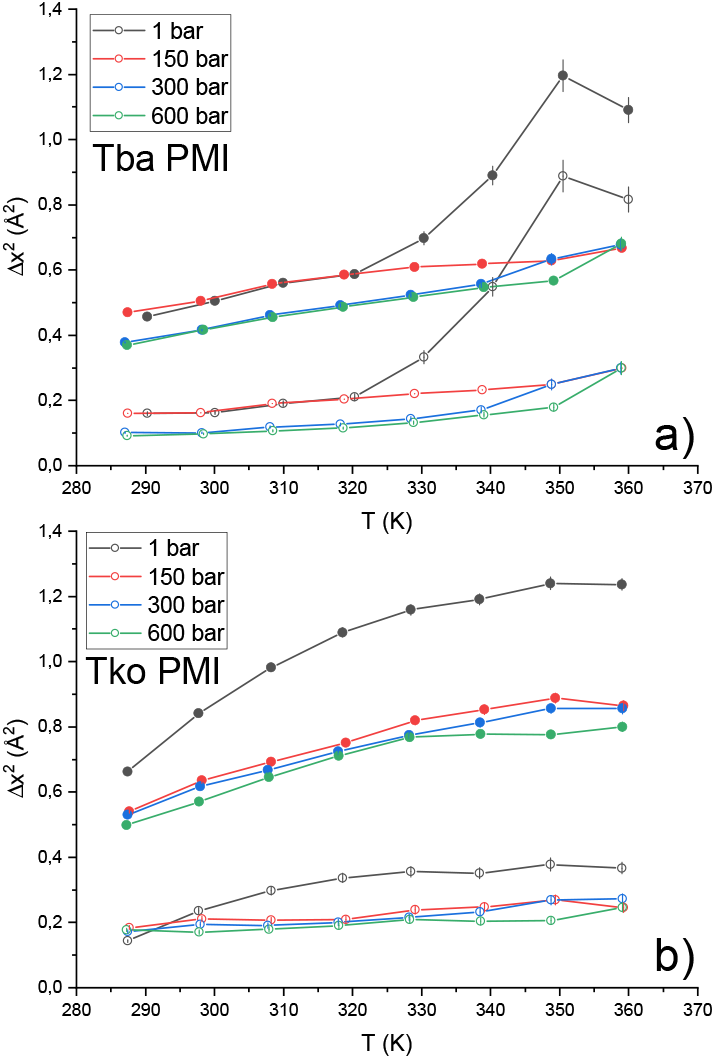
Total MSD (full circles) and MSD into the single wells (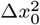, open circles) for Tba PMI (a) and Tko PMI (b). Lines are a guide to the eye.

**Figure 3:**
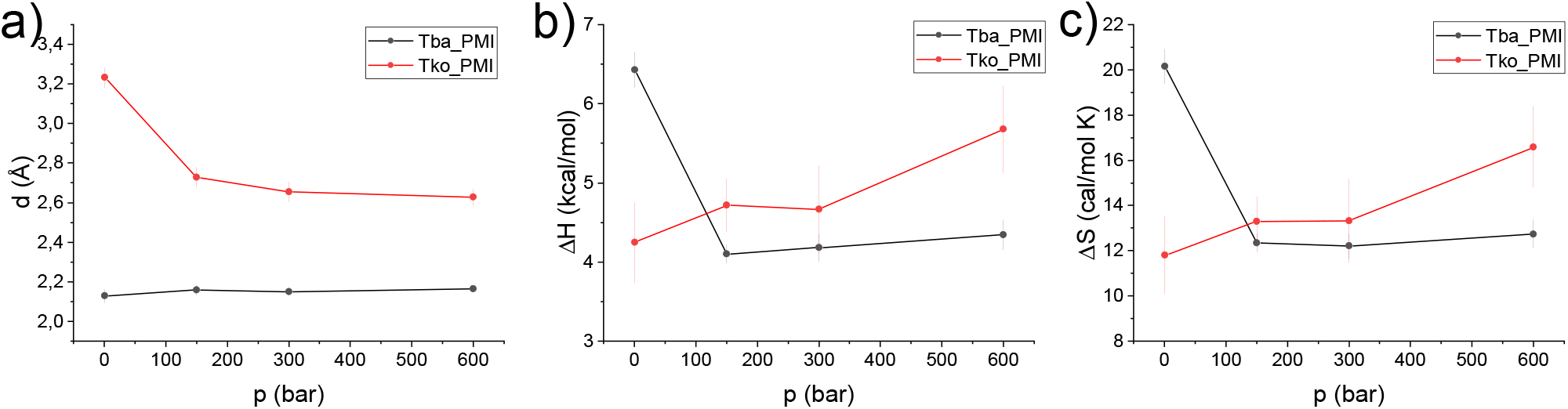
Temperature-independent parameters extracted from the two-state model as a function of pressure for Tba PMI (black symbols) and Tko PMI(red symbols): distance between the two wells (d, panel a), enthalpy (ΔH, panel b) and entropy (ΔS, panel c) difference. Lines are a guide to the eye.

### Quasi-elastic neutron scattering (QENS)

QENS was used to extract detailed information on the fast dynamics of the protein, to probe the effect of HHP, and to reveal how specific substitutions in the sequences could affect them, giving insight on the mechanism of pressure adaptation.

As in EINS, the QENS signal is dominated by the incoherent contribution of hydrogen atoms, but in this case a small energy transfer between the incident neutrons and the sample is allowed: it is thus possible to separate the contributions to the signal arising from different motions, while EINS gives an averaged picture. The analysis of the spectra gives access to localized and diffusional motions taking place on a specific time scale, which is ∼ 10 *ps* for IN5, and to characterize their geometry. Figure 4 shows a fit example, in which the two components we identified are highlighted: a broad and *q*-independent contribution due to fast localized motions, and a narrow contribution arising from confined jump-diffusion of protein residues. The logarithm of the HWHM of the broad component (fig. 5a and b) follows an Arrhenius behaviour (Γ_*loc*_(*T*) = Γ_0_ exp(−*E*_*A*_*/RT*), where *E*_*A*_ is the activation energy) at all pressure values for both samples. Tba PMI shows enhanced pressure stability with respect to its piezosensitive counterpart concerning fast localized motions. The activation energy values suggest that the rotation of methyl groups is the dominant process from which this contribution arises (values ranging from 1.5 to 3.8 *kcal/mol* have been reported^38^). Moreover, the value of *E*_*A*_ for methyl rotations has been shown to decrease in efficiently packed hydrophobic environments^38^. Hence, our data indicate that the extent of compression of the protein hydrophobic core is larger in Tko PMI than in Tba PMI, presumably due to the presence of bigger cavities in the former^12^. Concerning the narrow component, the extracted parameters are the mean jump length ⟨*l*⟩ and the residence time *τ*, which represents the mean time between two successive jumps. The temperature dependence of ⟨*l*⟩ (fig. 5c and d) for Tba PMI is rather weak and does not change with pressure, testifying the structural stability of the protein in the whole temperature and pressure range studied. For Tko PMI, this quantity shows a similar behaviour at 1 *bar*, where the protein is expected to be functional, while higher pressures seem to have a destabilizing effect. While it would appear straightforward to compare this quantity to the distance between the wells *d* derived from EINS data (fig. 3a), it must be stressed that the latter results from *all* the internal motions that are activated on the 100 *ps* time scale, and thus gives an average representation of the protein’s energy landscape, while ⟨*l*⟩ refers to a particular motion, namely the jump diffusion of side chains, and it relates to a different time scale. Furthermore, for the sake of comparison with other works, a *pseudo*-diffusion coefficient related to internal dynamics can be calculated according to *D*_*pseudo*_ = ⟨*l*⟩^2^ */*2*τ* ^39^ (fig. S15). The difference in the temperature dependence of *τ* (fig. 5e and f) for the two proteins is remarkable: it follows the Arrhenius law (*τ* (*T*) = *τ*_0_ exp(*E*_*A*_*/RT*), note the sign reversal compared to before, as *τ* = *ħ/*Γ) for Tba PMI, while it follows the Vogel-Fulcher-Tamman (VFT) law 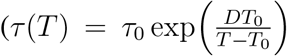, where 1*/D* is the fragility index, not to be confused with the aforementioned pseudo-diffusion coefficient *D*_*pseudo*_, and *T*_0_ is the Vogel temperature) for Tko PMI. The latter is typical of glass-forming systems^40–45^, but it has also been observed in proteins and interpreted as the signature of protein-water coupled dynamics^46–49^. An Arrhenius behaviour arises from activated processes, and it is expected for jump-diffusion, while a VFT behaviour is usually connected with co-operative processes. The appearance of VFT behaviour on such a fast time-scale is intriguing, and it shows that side-chain relaxations are strongly coupled to hydration water dynamics in Tko PMI. This correlates with the higher Δ*S* in the EINS data, and suggests that the decoupling of protein dynamics from its environment could be the key to pressure adaptation for Tba PMI. To push the analogy further, materials with a low fragility index exhibit a linear behaviour far from the Vogel Temperature *T*_0_ which can be reasonably fitted with the Arrhenius law, and are referred to as *strong* glass-forming materials, while *fragile* materials show VFT behaviour in a considerably wider temperature range^50^. Thus, the difference in dynamical properties between Tba PMI and Tko PMI could be assimilated to that between strong and fragile glass-forming materials. It appears that Tko PMI’s dynamics is dominated by cooperative motions, and that high pressure is able to destabilize them, as highlighted by the increase in fragility (i.e. decreasing *D*) with increasing pressure, while Tba PMI’s dynamics is dominated by pressure-insensitive activated processes. This difference is likely arising from the distinct amino-acidic composition of the two proteins, and it could explain the superior pressure stability of Tba PMI. Concerning the analysis of the EISF (fig. S16), the behaviour of the confinement radius *R* (fig. 5g and h) highlights another remarkable difference between the two proteins. Tba PMI shows a weak temperature dependence of this parameter, compatible with thermal expansion, and again shows no pressure dependence. On the other hand, the temperature dependence of Tko PMI appears to be stronger, and a sizeable increase of the confinement radius is detected with increasing pressure. This result could appear counter-intuitive, but it can be rationalized by thinking of *R* as an average measure of the protein’s solvent-accessible cavities: higher pressure forces water into them, increasing their volume, while concomitantly decreasing the protein specific volume, in agreement with Le Châtelier’s principle (fig. 6b). This volume increase in Tko PMI could also explain the enhanced mean jump length seen at high pressure (fig. 5d) and the stronger coupling with water displayed by the protein side-chains (fig. 3c and 5f).

**Figure 4:**
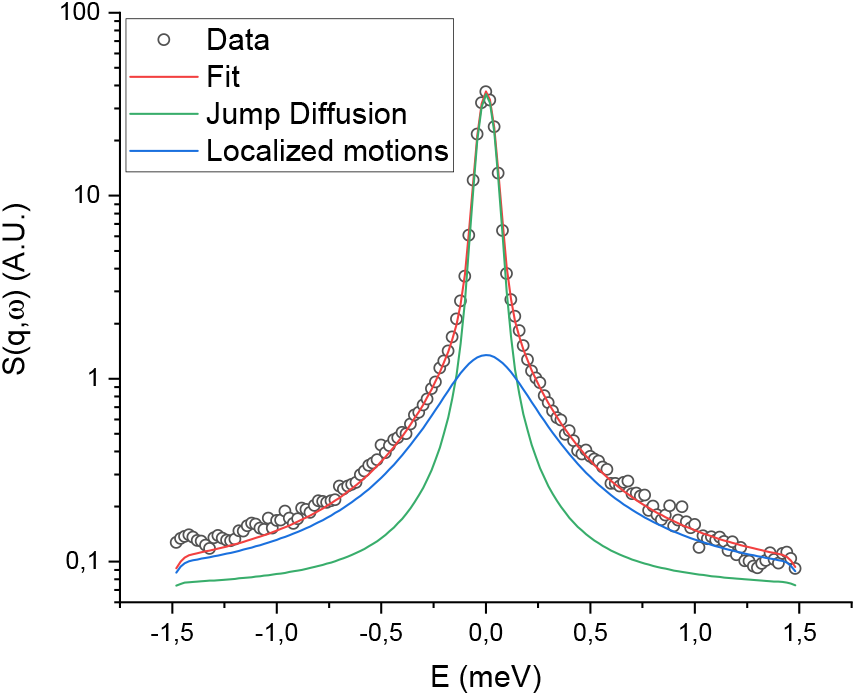
Fit example of Tba PMI at 1 bar and 286 K, at a q value of 1.14 Å^−1^. Black circles represent the corrected data, the two Lorentzian contributions owing to localized motions and jump-diffusion are shown respectively as blue and green solid lines, the total fit is shown as a red solid line. Error bars are smaller than the symbol size.

**Figure 5:**
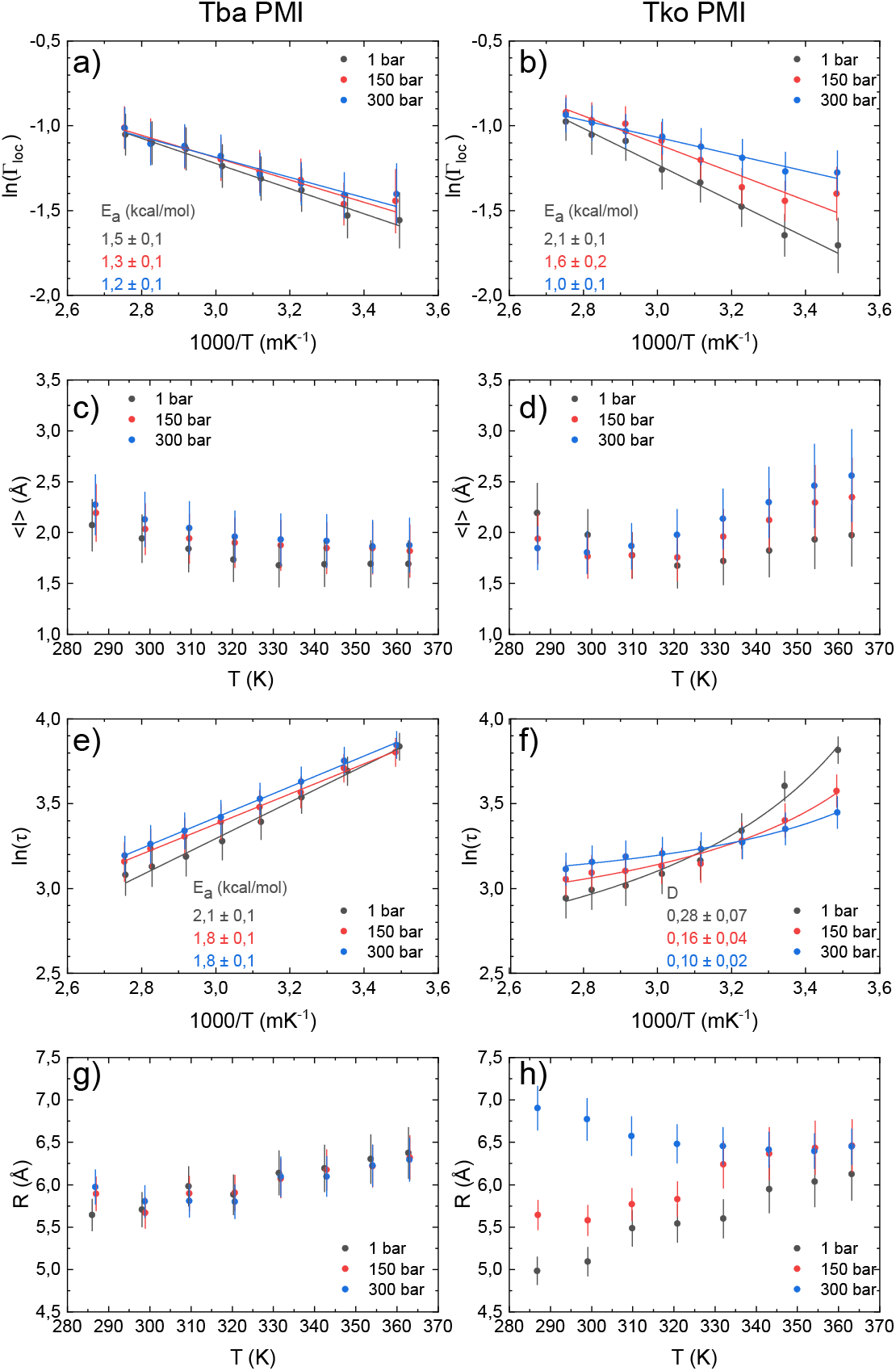
Natural logarithm of the broad component HWHM as a function of inverse temperature for Tba PMI (panel a) and Tko PMI (panel b) at all pressure values. Lines are linear fittings assuming an Arrhenius behaviour. Activation energy values are reported on the figure color-coded with the plots. Mean jump length from the Hall-Ross model as a function of temperature and pressure for Tba PMI (panel c) and Tko PMI (panel d). Natural logarithm of the residence time as a function of inverse temperature at all pressure values. Lines are fits to the Arrhenius law (panel e, Tba PMI) or to the Vogel-Fulcher-Tamman law (panel f, Tko PMI). Values for the activation energy (Tba PMI) and the fragility index (Tko PMI) are reported on the figure color-coded with the plots. Values of confinement radius R for Tba PMI (panel g) and Tko PMI (panel h).

**Figure 6:**
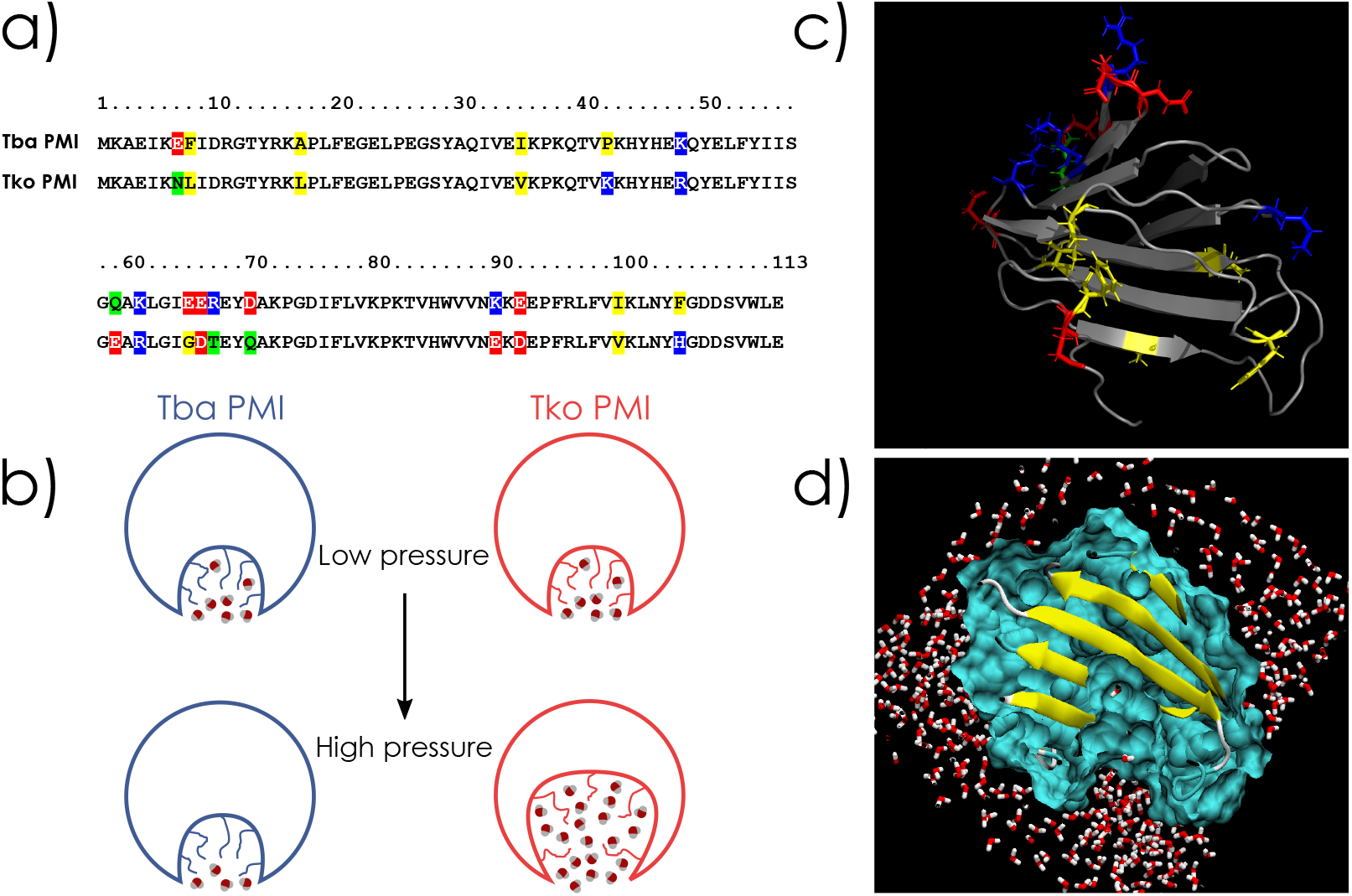
Panel a, sequence alignment of the two proteins, substitutions are highlighted (red for acidic, blue for basic, green for polar and yellow for hydrophobic residues). Panel b, schematic representation of the two proteins and effect of high pressure on them. Panel c, cartoon representation of Tba PMI, substitutions are represented in licorice with the same color-code as panel a. Panel d, vertical cut on the surface representation of the Tba PMI model, highlighting the ligand pocket.

## Discussion

The aim of this study is to compare proteins under positive selection originating from nearly isogenic organisms, differing only in their adaptation to pressure. Hence, for the first time, the different dynamical properties exhibited by the two proteins, as revealed by EINS and QENS, can be attributed to the difference in their sequence, reflecting the adaptation to HHP regardless of other adaptative traits. The results show that the mechanism by which Tba PMI counteracts the effect of HHP is by preventing excessive amounts of water from penetrating the solvent-accessible cavities and limiting their destabilization, as well as by inhibiting protein-water cooperative relaxations. This becomes of particular importance when considering the ligand pocket of the protein, i.e. the largest solvent-accessible cavity: its structure must be preserved under HHP in order to properly carry out the enzymatic reaction (fig. 6c) and to play its role in the metabolism of the organism. In view of these results, it is possible to characterize the impact of each substitution by comparing the two protein models and their sequences. The substitutions will be identified with the first letter being the residue present in Tba PMI, and the second letter for that in Tko PMI. The two proteins differ at 16 positions, shown on the 3D models of the protein (fig. 6a and b): half of the substitutions are present on the outer surface of the protein, 3 are at the entrance of the ligand pocket, 2 in the dimerization region and 2 near the putative active site. It is known that even small volume changes in the interior of a protein (such as a single point mutation) can greatly affect its pressure stability, especially when this residue is located close to a cavity and the substitution is affecting its volume^12^. To this extent, the I35V substitution appears to be key in the stabilization of the hydrophobic core, with the added importance of being close to the putative active site. The latter is very conserved except for K48R, which is not affecting charge or volume, and P42K, where the proline residue might soften the rigid *β*-strand structure and render it more resilient to pressure changes in Tba PMI. In the dimer-forming region, the A17L and I100V substitutions might optimize the contact between the two monomers, and decrease cavity volume. E7N, F8L and F105H, which are located at the ligand pocket entrance, appear to be the main substitutions that explain the observed difference in dynamics: indeed, the increased steric hindrance brought about by the phenylalanines together with the overall charge reversal might decrease water penetration into the pocket (fig. 6c and d) in Tba PMI. Lastly, the remaining substitutions (Q59E, K61R, E65G, E66D, R67T, D70Q, K90E, E92D), which are located at the protein-water interface, are likely involved in the modulation of the protein’s surface charge distribution, affecting its coupling with hydration water. Hence, the amino-acid substitutions between the two PMIs allow to draw a first adaptation pattern related to HHP. This is characterized by a decrease in polar residues and an increase in acidic and hydrophobic ones, and an enrichment in glutamate, lysine and phenylalanine in the piezophilic protein. It is also interesting to note that half of the substitutions affect the surface of the protein, which is consistent with the observations of an altered protein-water interaction. However, the relative contributions of the different amino-acid substitutions identified in this work need to be confirmed by further studies using and direct methods (multidimensional NMR spectroscopy, molecular dynamics simulations, site-directed mutagenesis), and eventually other piezophilic proteins.

## Acknowledgements

The authors would like to acknowledge the Institut Laue-Langevin for the allocation of beamtime to perform these experiments, James Maurice and the whole High-Pressure Division of the SANE group for technical support during the experiments, and Miguel Angel Gonzalez for his help during the re-writing of data reduction routines for LAMP. This work was supported by the French National Research Agency (programme ANR 17-CE11-0012-01 to P.O. and J.P.) and by the Mission pour les Initiatives Transverses et Interdisciplinaires of the CNRS (OriginsUnderPressure research project). A.C. is supported by a PhD grant for international students by the French Ministry of Science and Technology.

## Data Availability

Data are available at http://dx.doi.org/10.5291/ILL-DATA.8-04-876 for the IN13 experiment (8-04-876), and at http://dx.doi.org/10.5291/ILL-DATA.8-05-458 for the IN5 experiment (8-05-458).

## Methods

### Protein expression and purification

Recombinant Phosphomannose Isomerases from *T. barophilus* and *T. kodakarensis* have been produced by cloning synthetic codon-optimized genes (purchased from GENEWIZ Europe) into the protein over-expression plasmid pET-16b^24^ (Novagen), which was then transformed into *E. coli* BL21(DE3) pLysS strain (Novagen). 10 L cultures were grown at 37°*C* in LB medium supplemented with 100 *μ*g/ml ampicillin until *OD*_600_ = 0.5, induced with a final concentration of 1 mM IPTG and further grown overnight at 25°*C*. Cells were harvested by centrifugation at 17.000 g for 30 min, washed in isotonic solution (0.9% NaCl) and resuspended in 50 mM NaH_2_PO_4_, 300 mM NaCl, pH 8 buffer. Cells were then lysed by 5 freeze-thaw cycles in liquid nitrogen (1 min) and at 50°*C* (3 min), and homogenized by sonication (maximum power for 15 min at 50% duty cycle). The soluble fraction was recovered by centrifugation at 12.000 g and 4°*C* for 60 min. It was then heated at 75°*C* for 1 *h* to remove the non-thermostable proteins from the *E. coli* expression host. Protein debris were removed by centrifugation at 12.000 g and 4°*C* for 60 min. The supernatant was concentrated by ammonium sulfate precipitation and further purified by Size Exclusion Chromatography on an AKTA^®^ FPLC system, using an XK50-60 column packed with 1 L of Superdex^®^ 75 Prep-Grade resin, calibrated with the GE Healthcare^®^ Low Molecular Weight kit (fig. S2). During this step, both proteins evidenced a dimeric quaternary structure, as they eluted at double the expected MW (fig. S3). Fractions containing the protein were then pooled, concentrated by ultrafiltration (Amicon^®^ Ultra-15 centrifugal filter units, Millipore) and lyophilised. The purity of the proteins was assessed by SDS-PAGE, and was greater than 99% (fig. S4). To prepare the samples, the lyophilised protein powder was gently dissolved in D_2_O (Sigma-Aldrich) under nitrogen atmosphere, at a concentration of 120 *mg/ml*. Protein solutions rather than hydrated powders were employed in order to optimally transmit hydrostatic pressure to the sample. Proteins employed in the two different experiments belonged to the same production batch.

### Elastic incoherent neutron scattering (EINS)

EINS measurements were performed on the IN13 backscattering spectrometer at the Institut Laue-Langevin (ILL, Grenoble, France). At the elastic position, IN13 has an incident wavelength of 2.23 Å and a nearly *q*-independent resolution of 8 *μeV* FWHM, which gives a time window of ∼ 100 *ps*^51^, allowing to probe local motions of hydrogen atoms since their incoherent scattering cross section is an order of magnitude larger than that of other isotopes^30^. Temperature was controlled by means of a closed-cycle dry cryofurnace (Displex+), and continuous up-scans were performed in the 283 *K* to 363 *K* range at 0.08 *K/min*. The scattering intensity was also measured while the temperature was lowered back to 283 *K* before the next pressure point to check for hysteresis and, once its absence was verified, the downscans were merged with the upscans to improve statistics. HHP was transmitted to the sample by means of the high-pressure stick, cell and controller developed by the SANE team at ILL^52^, and four pressure points were investigated (1, 150, 300 and 600 *bar*). The high-pressure cell is cylindrical and made of a high-tensile aluminium alloy (7026) and has a 6 *mm* internal diameter^53^. A piston separates the pressure-transmitting liquid (Fluorinert^™^ FC-770^54^) from the sample, and a cylindrical aluminium insert (4 *mm* diameter) was used to decrease sample volume and to minimize multiple scattering. Raw data were corrected for transmission, empty cell and D_2_O scattering, normalized to a vanadium standard and then binned in temperature in 10 *K* intervals using the LAMP^55^ software available at ILL. EINS data have been interpreted in the framework of the *two-state model*,^32^ which models hydrogen atoms motions as a combination of vibrations in two harmonic potential wells, which give the Debye-Waller contribution with the associated Mean Square Displacement (MSD) 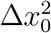, and jumps between them. The wells are separated by a distance *d* and have a free-energy difference Δ*G* that can be separated into the enthalpic and enthropic contributions according to Δ*G* = Δ*H* − *T*Δ*S*. The elastic scattering function *S*(*q, ω* = 0) as a function of the scattering vector *q* (related to the scattering angle *θ* and the neutron’s wavelength *λ* according to 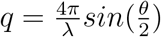) thus reads:

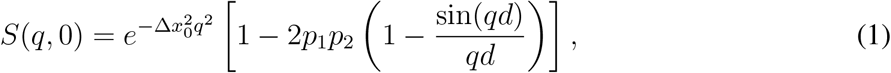

where *p*_1_ and *p*_2_ represent the population of each well, assumed in our case to follow the Arrhenius law (*p*_1_*/p*_2_ = *exp*(−Δ*H/RT* + Δ*S/R*), where *R* is the gas constant). It must be stressed, however, that in the investigated temperature range, large-scale motions could enter the experimental window. It is thus desirable to view the two wells as an average representation of the protein’s free-energy landscape that is accessible at each temperature and pressure value.

### Quasi-elastic neutron scattering (QENS)

QENS measurements were carried out on the IN5 time-of-flight (TOF) spectrometer^56^ at ILL at 5 Å incident wavelength. In this configuration the energy resolution was ∼ 70 *μeV HWHM*, giving a time window of ∼ 10 *ps*, suitable to investigate fast localized protein motions. Temperature was controlled with the standard ILL Orange Cryofurnace in the same range as the EINS experiment, and continuous scans at 0.4 *K/min* were acquired. The same HHP equipment was used for pressure transmission at 1, 150 and 300 *bar*. The 600 *bar* point could not be measured because of time constraints. The same corrections as in the EINS data treatment were applied (see SI) and, after temperature binning, TOF data were further corrected for detector efficiency and detailed balance^29^, then converted to *S*(*q, ω*) (where *ħω* is the energy that a neutron exchanges with the sample) and re-binned in 20 spectra with evenly spaced (0.02 *meV*) energy points at *q* values from 0.07 to 2.57 Å^−1^. Only spectra having a sufficient dynamic range (−1.5 to +1.5 *meV*) were considered in the analysis, giving a final *q* range of 0.6 − 1.8 Å^−1^. The whole treatment was performed with LAMP^55^. First, a model-free analysis of the corrected data was performed. This consists in fitting a sum of Lorentzian functions^57^ and leaving their parameters free in order to identify the different dynamical contributions to the measured signal, and then analysing the *q* dependence of their HWHM to define a suitable model that properly fits the data (see SI). Two main contributions have been identified in our case (adding a third Lorentzian did not improve the quality of the fit): the broad component displayed a substantially *q*-independent width, thus representing fast localized motions (e.g. methyl group rotations^31^), while the narrow component’s width exhibited a saturation behaviour at high *q*, characteristic of jump-diffusion processes of protein side-chains^58^. Among the different models that have been tested^39,59,60^, the Hall and Ross model^39^ gave the most satisfactory results (fig. S14). Therefore, the model function has been built by considering an elastic fraction (represented by the *Elastic Incoherent Structure Factor*, or EISF, *A*_0_(*q*)) plus a q-independent Lorentzian, representing the localized motions^29^ (Γ_*loc*_), and then convoluted by another Lorentzian which represents the jump-diffusion process in the Hall-Ross model^39^. This component is characterized by its *q*-dependent HWHM (Γ_*j*_(*q*)), which depends on the time between two successive jumps (*τ*, also named *residence time*) and the average length by which hydrogen atoms jump (⟨*l*⟩). The theoretical scattering function *S*(*q, ω*) thus reads:

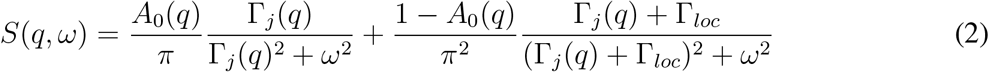

with

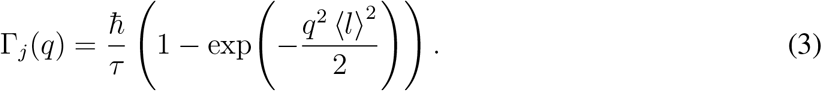

The model function is then convoluted with the resolution function (derived from a measurement of vanadium, as it is a dominant elastic incoherent scatterer, as shown in SI), multiplied by a *q*-dependent scale factor proportional to the Debye-Waller factor^29^, and then fitted to the data using a global fitting approach (i.e. by fitting the whole *S*(*q, ω*) at once instead of fitting single spectra at different *q* values separately), which gives the parameters Γ_*loc*_, *τ* and ⟨*l*⟩. In order to minimize the number of free parameters and to avoid ambiguities in the global fitting procedure, *A*_0_(*q*) has been calculated by integrating the spectra in the elastic region, and dividing this value by the total integral of the spectra, following its definition^29^. The calculated *A*_0_(*q*) have then been used as fixed parameters in the global fitting, permitting to fit the whole *S*(*q, ω*) with only three free parameters (Γ_*loc*_, *τ* and ⟨*l*⟩) and giving solid and consistent results. Global diffusion of the protein was not taken into account as the broadening arising from it would be lower than the resolution of the instrument in this configuration (see SI). To complete the picture, the geometry of these motions has been characterised by analysing the EISF. It has been modelled taking into account methyl rotation (*A*_3−*j*_ with 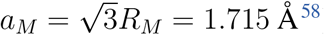) and restricted jump-diffusion of protein residues (*A*_*j*_, from the Hall-Ross model^39^) according to:

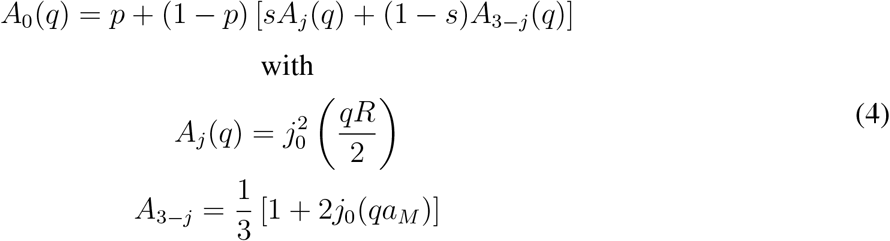

where *p* represents the fraction of immobile H atoms (i.e. slower than the time-scale of the experiment), *s* is the fraction of H atoms experiencing confinement during their jump-diffusion motion, *j*_0_ is the zeroth-order Bessel function of the first kind and *R* is the confinement radius.

## Notes

https://doi.ill.fr/10.5291/ILL-DATA.8-04-876

https://doi.ill.fr/10.5291/ILL-DATA.8-05-458

### Competing Interest Statement

The authors have declared no competing interest.

