## Supplementary Information for "Unravelling the mechanisms of adaptation to high pressure in proteins"

### Supplementary Informations

#### Genes and proteins sequences

The complete genomes of *Thermococcus barophilus* MP and *Thermococcus kodakarensis* KOD1 are available on GenBank under the accession codes CP002372[1] and AP006878[2] respectively. The two genes are identified as TERMP_00744 (Tba PMI) and TK_0503 (Tko PMI). The codon-optimized sequences used in this work to express the two proteins are reported here:

```
Tba PMI (TERMP_00744)
GCTGAACATATGAAAGCCGAAATTAAGGAGTTCATTGACAGGGGAACTTA
TAGGAAAGCCCCATTGTTTGAAGGTGAGCTTCCTGAAGGGAGTTACGCTC
AAATAGTTGAAATTAAACCCAAGCAGACGGTTCCAAAACACTATCATGAA
AAACAATATGAACTATTTTACATAATTAGCGGACAAGCAAAGCTCGGCAT
TGAAGAAAGAGAATATGATGCAAAACCAGGGGACATATTTTTAGTTAAGC
CCAAGACTGTTCATTGGGTTGTCAATAAAAAGGAAGAGCCATTCAGGCTT
TTTGTGATTAAGCTGAACTACTTTGGAGATGATAGCGTTTGGCTTGAGTG
AGGATCCTTCC
```

```
Tko PMI (TK_0503)
GGCTGAACATATGAAAGCAGAAATCAAAAATTTAATTGACCGCGGTACAT
ATCGTAAATTACCGTTATTCGAAGGTGAATTGCCTGAAGGTTCATACGCT
CAAATTGTAGAAGTAAAACCTAAGCAGACTGTAAAAAAACACTACCACGA
ACGTCAATACGAACTGTTCTACATTATCTCTGGTGAAGCTCGTCTTGGTA
TTGGTGATACAGAATATCAAGCAAAGCCTGGTGATATTTTTCTAGTTAAA
CCAAAAACCGTACACTGGGTAGTTAATGAAAAAGACGAACCATTCCGTCT
TTTTGTTGTTAAGTTAAATTATCACGGTGATGACTCTGTATGGTTAGAAT
GAGGATCCTTCC
```

The restriction sites for *NdeI* (blue) and *BamHI* (red) have been indicated. The genes were received from GENEWIZ in pUC57-Kan plasmid, digested with *NdeI* and *BamHI*-HF® (New England Biolabs) and then ligated into the Multi Cloning Site of the pET-16b plasmid (digested with the same enzymes) by means of T4 DNA Ligase (Thermo Fisher). All construct propagation was done using NEB-5$\alpha$® *E. coli* cells (New England Biolabs). Plasmids were then extracted, purified (Macherey-Nagel NucleoSpin® Plasmid kit) and sequenced to verify the constructs, then transformed into the BL21(DE3) pLysS strain for expression. All the listed products have been employed according to the standard protocols issued by the manufacturers.

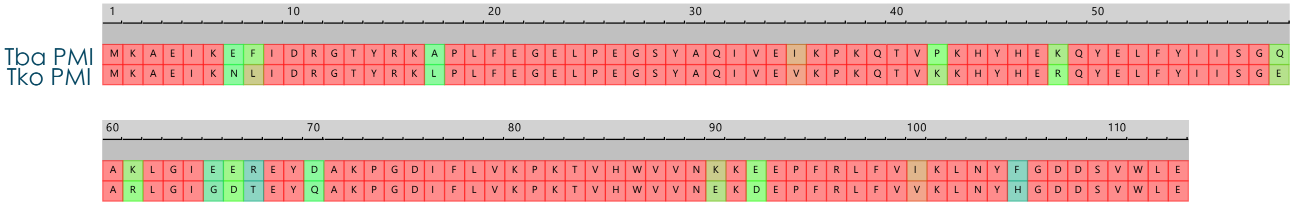


*Figure S 1: Sequence alignment of Tba PMI and Tko PMI, color coded for sequence similarity in a Blue-Green-Red scale (blue for least similar, red for most similar).*

Figure S1 reports the aligned sequences of the two proteins. They both are 113 residues long, and there are 16 substitutions between them, which are color-coded for sequence similarity (BLOSUM30) in a Blue-Green-Red scale (blue being least similar, red most similar).

#### Protein production and purification

The proteins have been expressed and purified as reported in the Methods section in the main text. Here we report the calibration run for the XK50-60 column using the GE Healthcare® Low Molecular Weight kit (fig. S2), which consists of five proteins: Conalbumin (75 kDa), Ovalbumin (44 kDa), Carbonic Anhydrase (29 kDa), Ribonuclease A (13.7 kDa) and Aproritin (6.5 kDa). The void volume of the column was determined to be $360\ ml$ by using the Blue Dextran 2000 supplied with the kit. The kit was used according to the manufacturer's protocol.

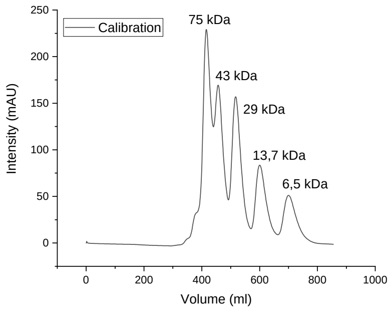


*Figure S 2: FPLC calibration run.*

Figure S3 reports an example purification run for Tko PMI, showing that it elutes at a volume compatible with a MW of 30 kDa (to be compared with fig. S2), but SDS-PAGE analysis of the fractions shows a band at the 15 kDa mark, thus confirming the dimeric nature of the protein.

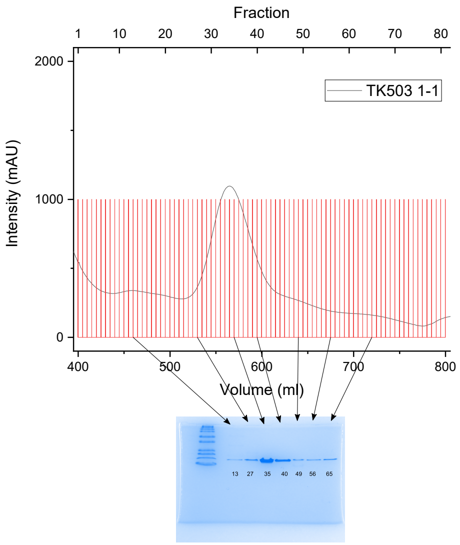


*Figure S 3: Example FPLC run. Fractions are given on the upper axis, and the SDS-PAGE analysis of some representative ones is given.*

Figure S4 reports an SDS-PAGE analysis of Tba PMI after pooling of the FPLC fractions and concentration, attesting the purity of the protein. The only steps performed after this check were lyophilization, and then the subsequent dissolution in $D_2O$ immediately before the neutron experiments.

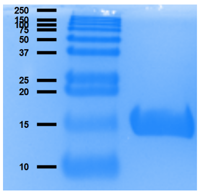


*Figure S 4: SDS-PAGE analysis of the final product (Tba PMI).*

#### Homology modelling and diffusion coefficient calculation

Homology modelling was performed on two different servers: i-Tasser[3] and Swiss-Model[4]. Both gave very similar models (fig. S5) and both correctly predicted the protein to have a dimeric quaternary structure (fig. S6).

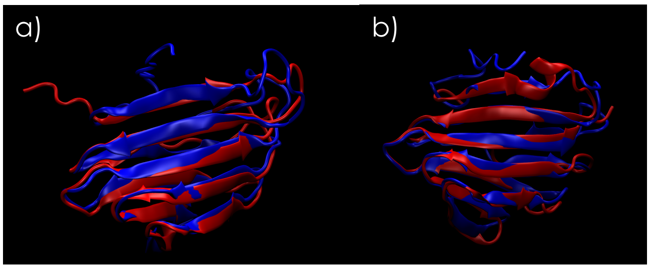


*Figure S 5: Structural alignment of the two models from i-Tasser (blue) and Swiss-Model (red) for Tba PMI (panel a) and Tko PMI (panel b).*

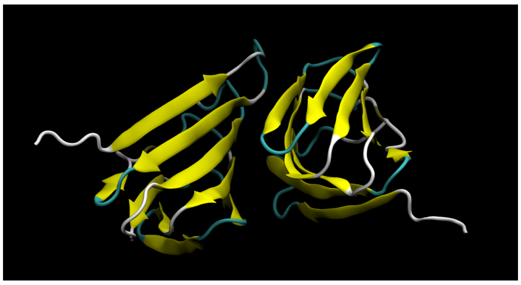

*Figure S 6: Dimeric model predicted by Swiss-Model for Tba PMI.*

Figure S7 shows the putative active site, located into the $\beta$-barrel and modelled as a manganese ion coordinated by His44, His46, Glu51 and His85. These residues are conserved between the two organisms.

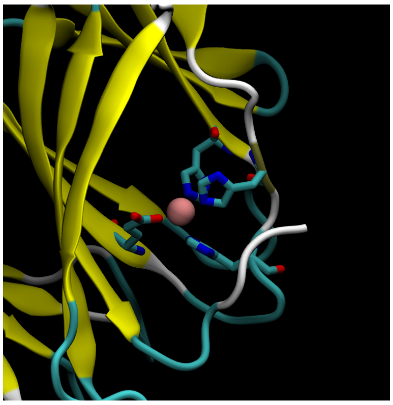

*Figure S 7: Detail of the ligand binding site for Tba PMI. The involved residues (His44, His46, Glu51 and His85) and the putative metal ion (Mn) are shown.*

Figure S8 shows the substitutions in solvent-exposed residues (Q59E, K61R, E65G, E66D, R67T, D70Q, K90E, E92D), colour-coded for residue type (blue for basic, red for acidic, green for polar), and figure S9 displays the ligand pocket entrance for both proteins after structural alignment. It is clear how the substitutions affect the available area for the solvent to enter the pocket. For a deeper discussion about the impact of the substitutions on the structure and dynamics of the two proteins, see the main text.

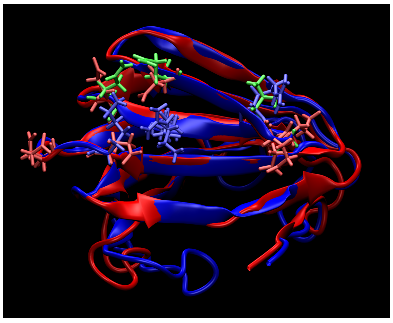

*Figure S 8: Structural alignment of Tba PMI (blue) and Tko PMI (red) highlighting the substitutions of solvent-exposed residues (blue for basic, red for acidic, green for polar).*

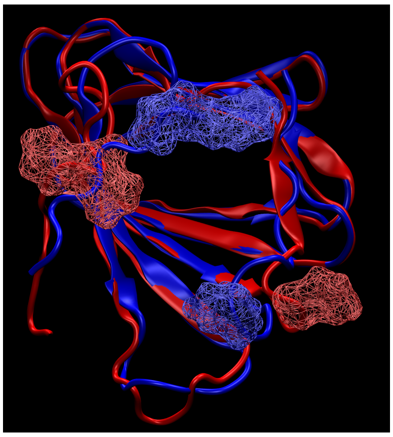

*Figure S 9: Detail of the ligand pocket entrance showing the residues Glu7, Phe8 and Phe105 for Tba PMI (blue), and Asn7, Leu8 and His105 for Tko PMI (red) in surface representation.*

The diffusion coefficient was calculated with HYDROPRO[5] for both proteins at 353 K (at atmospheric pressure). This was the highest temperature (and lowest pressure) reached during experiments, the result would thus give an upper limit for $D$. Moreover, HYDROPRO calculates $D$ in the infinite dilution limit, therefore we expect the real diffusion coefficient in our samples at 120 mg/ml to be considerably lower. The calculation was performed for both proteins by using the Swiss-Model dimeric model, and by using both the density and viscosity of pure $D_2O$. The results were $14.7\ \mathring{A}^2/ns$ and $15.1\ \mathring{A}^2/ns$ for Tba PMI and Tko PMI respectively, thus giving a quasi-elastic broadening of $\Gamma = \hbar D q^2 = 25.3 \mu eV$ for Tba PMI, and $27.4 \mu eV$ for Tko PMI at the highest $q$ value investigated here. Comparing this with the resolution of IN5 in our experimental conditions ($70\ \mu eV$ HWHM), we can conclude that translational diffusion of the proteins does not give a measurable contribution to the signal.

#### EINS and QENS Analysis

Both EINS and QENS data were corrected for empty cell, solvent scattering and detector efficiency (measured by means of a vanadium standard) by taking into account their transmission and the solvent volume fraction, according to the following expression:

$$S_{corr}(q,\omega) = \frac{(\frac{1}{t_{sample}} S_{sample} - \frac{1}{t_{cell}} S_{cell}) - \phi(\frac{1}{t_{D2O}} S_{D2O} - \frac{1}{t_{cell}} S_{cell})}{\frac{1}{t_{vana}} S_{vana}} \quad (1)$$

where $t$ denotes the transmission, and the $(q,\omega)$ dependence has been dropped for clarity. $\phi$ denotes the solvent volume fraction in the samples, which has been calculated by dividing the elastic scattering intensity of the solvent by that of the sample at around $q = 1.6\mathring{A}^{-1}$ at each temperature and pressure value, where $D_2O$ displays a broad coherent peak. The transmission can be directly measured on IN13 and the measured values were used for both data sets in virtue of the very weak dependence of the transmission on the neutron wavelength (2.23Å for IN13, 5Å for IN5). The transmission of the samples was 91% for Tba PMI and 93% for Tko PMI, we could thus reasonably assume that multiple scattering effects are negligible. Figure S10 shows the temperature independence of $d$ when fitting the IN13 data with the two-state model[6], thus justifying the global fitting approach with $d$ as a shared parameter.

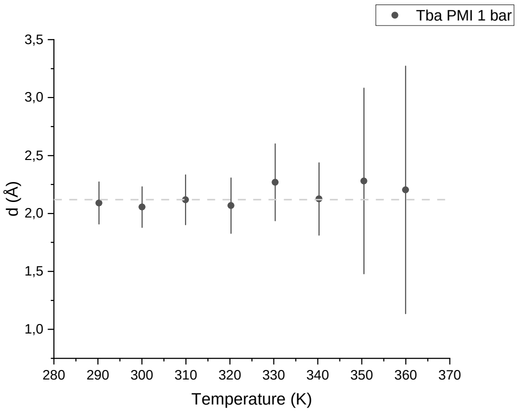


*Figure S 10: Temperature independence of the parameter d in the two state model.*

Figure S11 displays a vanadium spectrum, from which the resolution function was extracted by fitting a Gaussian curve. Its parameters at all $q$ values were then used to model the resolution function and convolute it to $S(q,\omega)$ to get the experimental scattering function $S_{exp}(q,\omega)$, according to:

$$S_{exp}(q,\omega) = B(q) + \mathcal{R}(q,\omega) \circledast [D(q)S(q,\omega)] \qquad (2)$$

where $B(q)$ is a flat background accounting for fast vibrational motions, $\mathcal{R}(q,\omega)$ is the resolution function, $\circledast$ is the convolution operator in $\omega$, $D(q)$ is a scale factor proportional to the Debye-Waller factor and $S(q,\omega)$ is the theoretical scattering function.

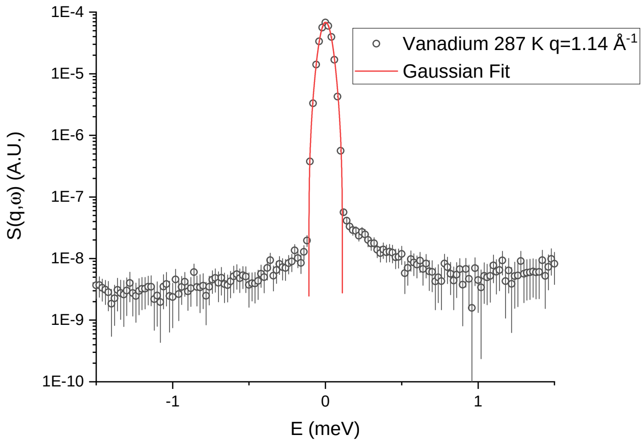


*Figure S 11: Spectrum of vanadium at 287 K and at* $q = 1.14\mathring{A}^{-1}$ *(black circles) and gaussian fitting (red line).*

We first performed a model-free approach by fitting a sum of Lorentzian functions and leaving their parameters free. As shown in Figure S12, when fitting with three Lorentzians the third always converged to a negligible contribution: its area was two orders of magnitude lower than that of $\mathcal{L}_1$ and $\mathcal{L}_2$, and its width was higher than the instrument's dynamic range in our conditions, therefore two Lorentzians were used. Their HWHM ($\Gamma$) are shown in Figure S13 and S14.

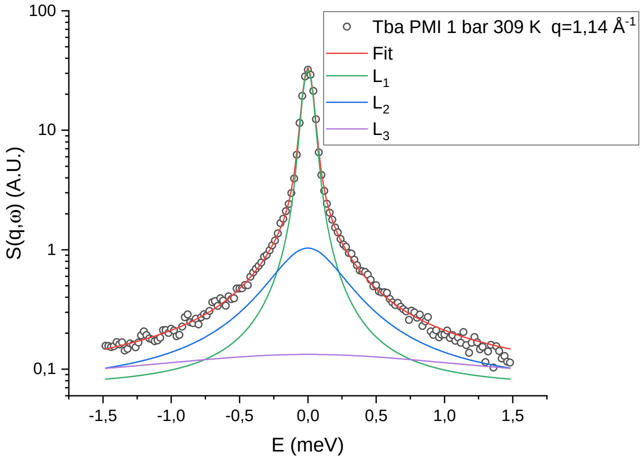


*Figure S 12: Example of a three-Lorentzians fitting.*

The broad component $\mathcal{L}_2$ shows a rather weak $q$-dependence (fig. S13), it has thus been assigned to localized motions, which give rise to a $q$-independent Lorentzian plus an elastic contribution[7], according to:

$$S_{loc}(q,\omega) = A_0(q)\delta(\omega) + \frac{1 - A_0(q)}{\pi} \frac{\Gamma_{loc}}{\Gamma_{loc}^2 + \omega^2} \tag{3}$$

where $A_0(q)$ is the *Elastic Incoherent Structure Factor*, and $\Gamma_{loc}$ is the $q$-independent HWHM related to localized motions.

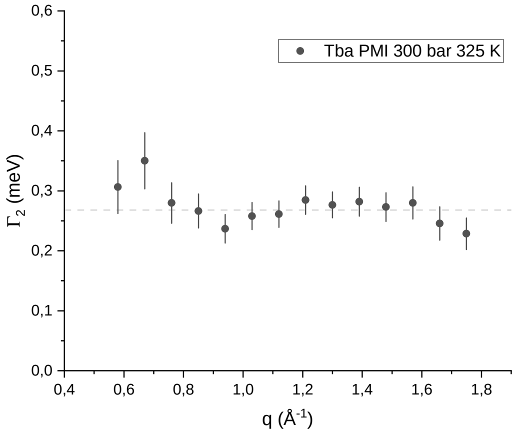


*Figure S 13: Width of the broad component as a function of the scattering vector q.*

The width of the narrow component $\mathcal{L}_1$ exhibited the signature characteristics of jump-diffusion motions, that is, monotonically increasing at low $q$ and reaching a plateau value at high $q$ (fig. S14). Different jump-diffusion models have been tested[8–10], and the one by Hall and Ross[8] was found to better fit the data compared to the one by Singwi and Sjölander, commonly used to model jump-diffusion motions in proteins.

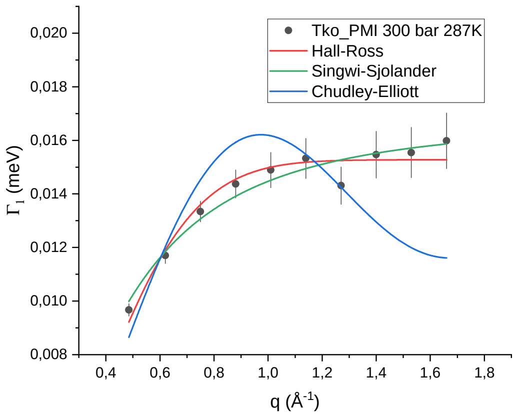


*Figure S 14: Width of the narrow component as a function of the scattering vector q.*

These motions thus bring about another Lorentzian contribution, according to:

$$S_{jump}(q,\omega) = \frac{1}{\pi}\frac{\Gamma_j(q)}{\Gamma_j^2(q)+\omega^2} \tag{4}$$

with

$$\Gamma_j(q) = \frac{\hbar}{\tau}\left(1-\exp\left(-\frac{q^2\left\langle l\right\rangle^2}{2}\right)\right). \tag{5}$$

where $\tau$ is the mean time between two jumps (residence time) and $\langle l \rangle$ is the mean jump length, that is, the mean of the jump distribution function, which is assumed to be Gaussian according to the Hall-Ross model. The overall theoretical scattering function (eq. 2 in the main text) results from the convolution of these two contributions. Since hydrogen atoms motions dominate the QENS signal, a convolution is necessary instead of a simple sum, as they can concomitantly perform both motions. As an example, a methyl group in an isoleucine can rotate (giving rise to $S_{loc}$) and, at the same time, perform jump-diffusion with the rest of the residue's side-chain (giving rise to $S_{jump}$).

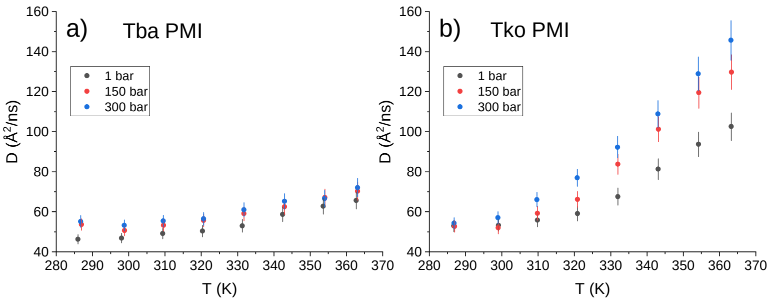


*Figure S 15: Pseudo-diffusion coefficient calculated for Tba PMI and Tko PMI at all temperature and pressure values.*

Figure S15 shows the pseudo-diffusion coefficient $D_{pseudo}$ described in the main text, which can be used to compare these results with other works.

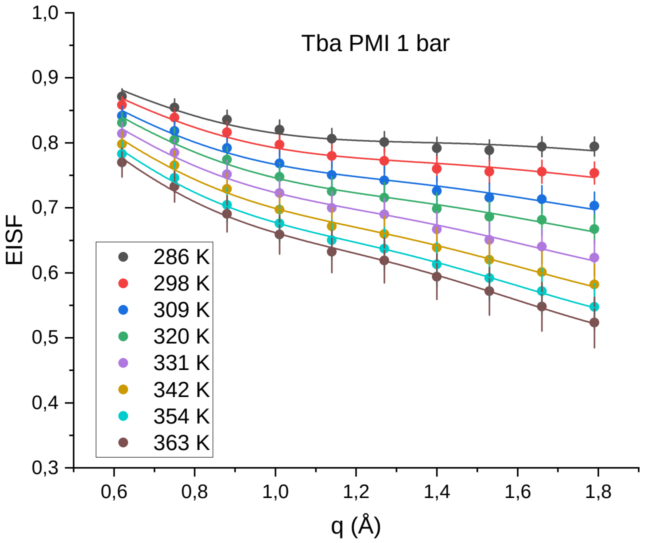


*Figure S 16: Example fit of the EISF at all temperatures for Tba PMI at 1 bar.*

Figure S16 shows a fit of the EISF at all temperatures for Tba PMI at 1 bar, according to the model defined in equation 4 in the main text, and figure S17 displays the fitting parameters not shown in the main text.

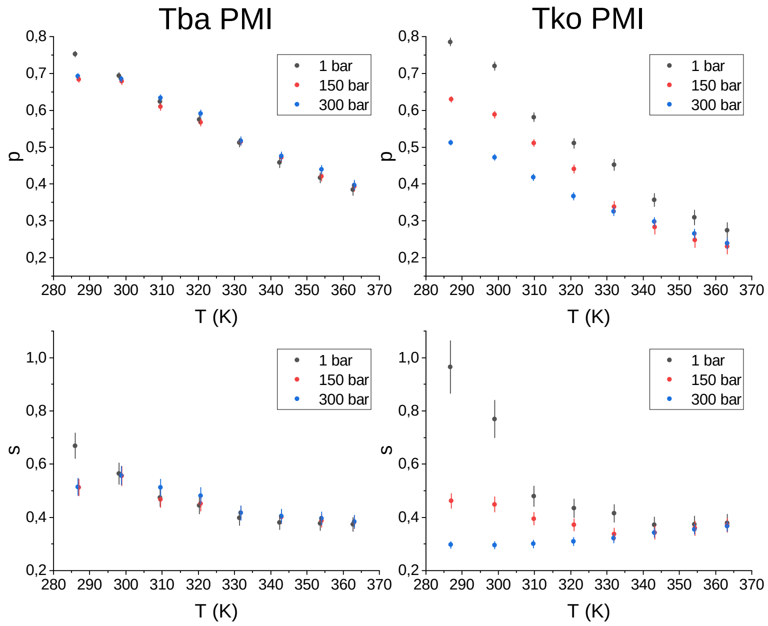


*Figure S 17: Temperature independence of the parameter d in the two state model.*

Both $p$ (immobile fraction) and $s$ (fraction of atoms performing confined jump-diffusion) are essentially pressure-independent, in line with all the other results, while for Tko PMI it is evident that pressure is activating new motions on one hand (decrease in $p$), and inhibiting the jump-diffusion motions on the other hand.
